# Long-range coupling regulates stator dynamics in the bacterial flagellar motor

**DOI:** 10.1101/2025.09.28.679083

**Authors:** Shabduli A. Sawant, I. Can Kazan, Brennen M. Wise, S. Banu Ozkan, Navish Wadhwa

## Abstract

The bacterial flagellar motor generates torque through MotAB stator complexes which couple ion flux to rotation. Stators anchor in the peptidoglycan cell wall and dynamically remodel in response to changes in external conditions such as the mechanical load, yet how stator anchoring is regulated remains unknown. Here, we show that long-range allosteric interactions within the MotB periplasmic domain tune stator binding in *Escherichia coli*. Using coarse-grained elastic-network modeling and co-evolutionary analyses, we identified residues mechanically coupled to peptidoglycan-interacting loops of MotB. Targeted mutagenesis at these coupled sites produced distinct motility phenotypes in some mutants, exhibiting altered swimming speeds compared to wild-type and characteristic expression-dependent swimming trends, indicating mutation-specific effects on stator dynamics or torque. Single-motor measurements distinguished mutants with altered torque from those with altered stator dynamics. Molecular dynamics simulations revealed that mutations at distal positions reshape loop flexibility in ways that quantitatively correlate with swimming speeds. These results demonstrate that allosteric communication within MotB propagates across length scales to modulate the performance of the entire motor, revealing how local molecular changes can tune large-scale bacterial motion.

## INTRODUCTION

Many bacteria use rotary flagella to navigate their environment, propelling themselves through complex and varied habitats (Wadhwa and Berg, 2022). Beyond simple locomotion, motility is often essential for infection, surface attachment, and biofilm formation (Haiko and Westerlund-Wikström, 2013; Belas, 2014). The ability to move across these distinct niches requires propulsion that is both robust and adaptable. Uncovering how bacteria generate and control this propulsion is therefore key to understanding the breadth of motility-driven behaviors.

The bacterial flagellum is a multi-component nanomachine composed of a helical filament, a flexible hook, and a rotary motor embedded in the cell envelope (Wadhwa and Berg, 2022; Sowa and Berry, 2008; Nakamura and Minamino, 2024). Rotation of this motor is powered by transmembrane stator complexes, each assembled from MotA and MotB in a 5:2 ratio (Stolz and Berg, 1991; Santiveri et al., 2020; Deme et al., 2020). Within each complex, the MotB dimer anchors the stator to the peptidoglycan and the P-ring from the flagellar bushing (De Mot and Vanderleyden, 1994; Roujeinikova, 2008; Hizukuri et al., 2010), while the MotA pentamer rotates around it to exert torque on the rotor (Hu et al., 2022) (Fig. 1a). Ion flux through the MotA–MotB complex provides the energy that drives this rotation (Meister et al., 1987). Collectively, these elements constitute the motor’s force-generating machinery, with MotB’s periplasmic domain (Fig. 1b) serving as the anchor that secures the stator to the cell wall.

**Figure 1.**
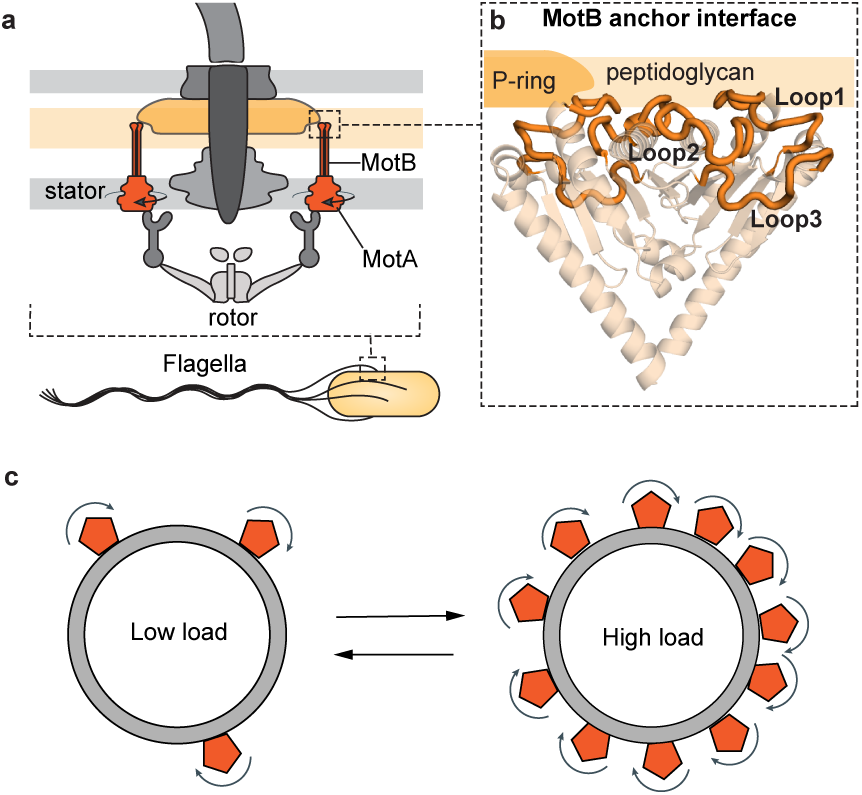
Load-dependent remodeling of torque-generating stators in the bacterial flagellar motor. **a** Key components of the motor, including the MotAB stator complexes (orange) that generate torque. MotA interacts with the rotor, while the periplasmic domain of MotB anchors the stator in the peptidoglycan near the P-ring (yellow). **b** Structure of the *E. coli* MotB periplasmic domain, obtained using SWISS-MODEL. Peptidoglycan interfacing loops are highlighted. **c** Load dependent remodeling of the stator binding. At low load, stators exchange rapidly between the motor and the membrane pool. At high load, stator anchoring is stabilized, reducing their off-rate and increasing the number of active stators.

Although the stator is anchored to the cell wall, its association with the motor can be highly dynamic. For instance, in *Escherichia coli* and *Salmonella enterica*, individual stator units continuously bind to and dissociate from the motor, allowing the complex to remodel in real time (Leake et al., 2006; Block and Berg, 1984; Blair and Berg, 1988). This turnover is strongly load dependent (Fig. 1c): stators dissociate rapidly under low load but remain bound when the motor experiences higher external load, a behavior characteristic of a catch bond (Lele et al., 2013; Tipping et al., 2013; Nord et al., 2017; Wadhwa et al., 2019, 2021, 2022). As a result, motors typically operate with around five bound stators but can recruit up to eleven under high load (Reid et al., 2006; Niu et al., 2023). Yet how external load, applied far from the MotB periplasmic domain, tunes its anchoring to the cell wall remains poorly understood.

Conformational changes in MotB, including linker opening and periplasmic-domain dimerization, play key roles in stator activation and anchoring (Kojima et al., 2009; Zhu et al., 2014; Kojima et al., 2018). This domain features three surface-exposed loops, 1, 2, and 3, positioned to interact with the surrounding periplasmic environment (Fig. 1b). Together with adjoining helices, the loops shape the surface that mediates MotB’s anchoring to the peptidoglycan and the flagellar P-ring (Hizukuri et al., 2010). Crystal structures of *Helicobacter pylori* MotB reveal that truncations in the N-terminal linker region produce large changes in loop conformation, suggesting that linker conformation exerts long-range control over loop positioning (O’Neill et al., 2011). Together with their linker-dependent conformational dynamics (Reboul et al., 2011), these observations raise the possibility that long-range allosteric interactions within the MotB periplasmic domain regulate loop conformation and, ultimately, stator anchoring.

Here, we combine structural modeling, molecular dynamics simulations, targeted mutagenesis, and quantitative motility assays to investigate how mechanical information is transmitted within the MotB periplasmic domain to influence stator function. We identify long-range dynamic coupling between distal residues and the peptidoglycan-binding loops, demonstrating that mutations at these sites differentially affect stator anchoring dynamics and torque output, and show that loop flexibility quantitatively correlates with swimming performance. Together, our results establish MotB as a mechanically integrated element that transmits force across nanometer scales via allosteric coupling, providing a physical mechanism by which bacteria tune motor output in response to external mechanical load.

## RESULTS

### Flexibility and long-range coupling in the MotB periplasmic domain

To investigate the intrinsic dynamics of the MotB periplasmic domain, we generated structural models of *E. coli* MotB using SWISS-MODEL (Waterhouse et al., 2018) and selected three representative conformations corresponding to distinct crystal forms observed in *Salmonella* MotB (Kojima et al., 2009). These models showed excellent agreement with AlphaFold3 predictions, with all three conformations superimposing onto the AlphaFold3 model with low RMSD (Fig. S1). For each model, we computed the Dynamic Flexibility Index (DFI), which quantifies the normalized displacement of each residue under random perturbations in an elastic network model (Gerek et al., 2013).

Across all three structural models, the DFI profiles revealed a consistent pattern: loop 3 and the H1 helix exhibited high flexibility, while loops 1 and 2 largely exhibited intermediate flexibility with some rigid portions (Fig. S2). In contrast, residues forming the core and the dimer interface were comparatively rigid (Fig. 2a). The agreement across the three models indicates that these dynamic features are robust to structural variability. Notably, the flexible loop 3 corresponds to the region known to interface with the P-ring in the intact motor (Hizukuri et al., 2010), suggesting that local mobility may play a role in tuning stator anchoring.

**Figure 2.**
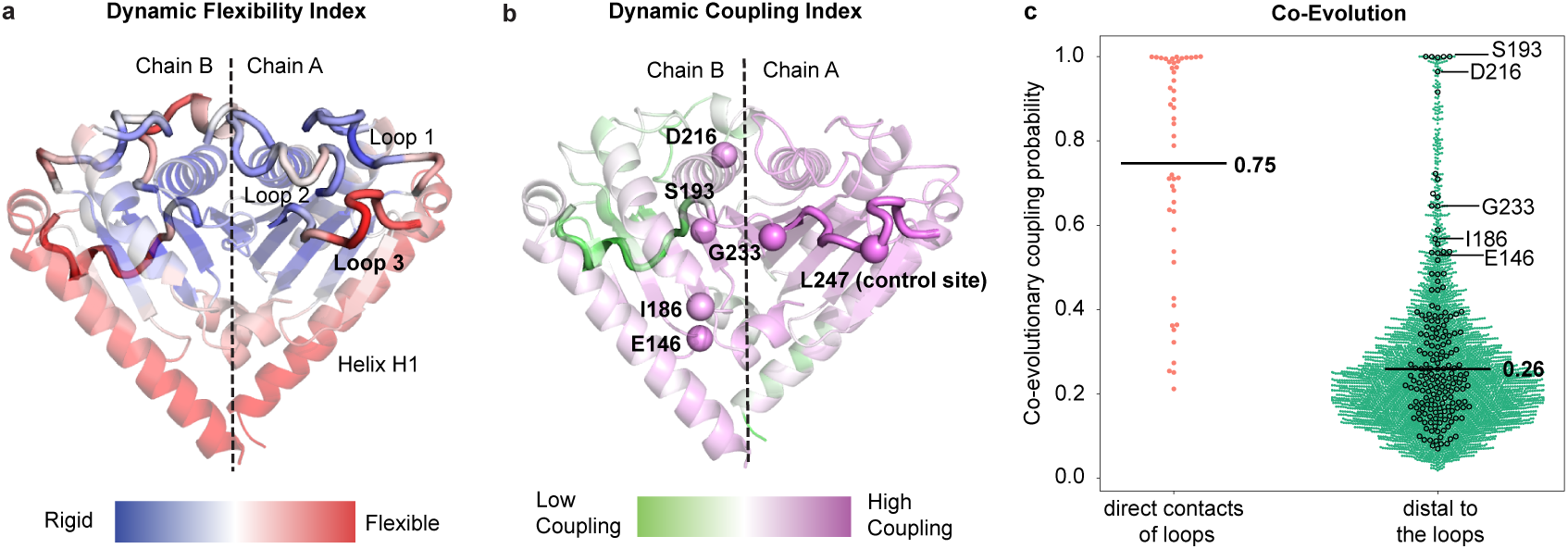
Dynamic and evolutionary couplings within the MotB periplasmic domain indicate conserved long-range interactions. **a** Flexibility profile of the MotB periplasmic domain. Loops 3 at the peptidoglycan interface, along with helix H1, display higher flexibility. **b** Dynamic coupling of chain A loop 3 with other regions of the MotB periplasmic domain. Residues in chain B that couple strongly with loop 3 in chain A are represented as spheres. Note that residue G233 represented as a sphere is part of chain B. **c** Dynamically coupled residues exhibit high coevolutionary coupling with loops 1, 2 and 3. Pairwise coevolutionary coupling probabilities are shown for residues in direct contact with the loops (<6.8 Å) and those that are distal to the loops (>8.0 Å). Average co-evolutionary coupling for each group is represented by the black horizontal line. Evolutionary coupling probabilities of residues from the high DCI group are plotted as black circles.

Because regulating anchoring requires transmitting load information from distal sites to the pepti-doglycan interface, we next used the Dynamic Coupling Index (DCI) (Ose et al., 2022) which uses the perturbation method (Kumar et al., 2023) to identify residues mechanically coupled to loop 3. Given loop 3’s high flexibility and its known interaction with the P-ring (Hizukuri et al., 2010), we designated it as the critical functional site for this analysis. Perturbing loop 3 revealed strong intra-chain coupling within the flanking helices and adjacent loop regions of the same monomer, consistent with local structural connectivity (Fig. 2b). Surprisingly, we also identified clusters of residues in the opposite monomer that showed high DCI values despite some of these being separated by 15–30 Å from loop 3 (Fig. 2b & Table S1). These regions included residues 146, 186, 193, 216, and 233, spanning both the dimer interface and distal elements of the domain. The presence of these long-range intra- and inter-chain couplings indicates that motions at loop 3 are mechanically linked to distant regions across the domain rather than being confined to the local environment.

### Coevolution supports long-range functional coupling in MotB periplasmic domain

To assess whether dynamically coupled residues also show signatures of co-evolving, we analyzed co-evolutionary couplings between MotB peptidoglycan interface and distal sites (Mishra et al., 2019) using EVcouplings, which infers residue–residue relationships from multiple-sequence alignments (Hopf et al., 2019). Because strong coevolution coupling scores is typically observed between residues in close physical proximity, high coupling probabilities between residues separated by more than ∼8 Å —the threshold for direct C*α* contacts (Green et al., 2021)—indicate long-range functional interactions. As expected, residues directly contacting loops 1–3 (<6.8 Å) exhibited higher average evolutionary coupling scores than distal residues (>8 Å) (Fig. 2c). However, distal residues that were strongly coupled to loop 3 in the DCI analysis showed unexpectedly high coevolution scores, approaching those of the direct-contact group (Fig. 2b). These overlaps indicate that long-range communication pathways within the MotB periplasmic domain are both mechanically and evolutionarily coupled Kazan et al. (2022). A summary of DFI, DCI, and coevolution results is provided in Table S1.

### Targeted mutations identify distal residues that modulate stator function

Guided by the DFI, DCI, and coevolution analyses, we selected candidate residues likely to participate in long-range regulation of the MotB periplasmic domain. To assess their functional roles *in vivo*, we introduced point mutations at two flexible positions (E146, I186), two rigid positions (S193, D216), one intermediate position (G233), and a loop-3 control site (L247). To avoid abolishing function entirely, amino acid substitutions were chosen based on ConSurf conservation scores (Fig. S3) with additional charge-altering mutations included to test the effect of more dramatic perturbations. These mutant stators were expressed from a pBAD33 plasmid in an *E. coli* strain lacking the native MotAB genes.

All mutants except the L247P active-site variant supported motility, confirming successful expression and assembly into functional stators. Motility assays revealed a wide range of phenotypes (Fig. 3a). Growth rates were comparable across all variants (Fig. S4), indicating that differences in motility were not attributable to differences in cell growth. Differences in soft-agar colony expansion reflected changes in stator function caused by these point mutations (Fig. 3a). These trends were corroborated by single-cell tracking in liquid, where swimming speeds paralleled the soft-agar expansion phenotypes (Fig. 3c), indicating that the motility differences arise from mutational perturbation of residues coupled to the peptidoglycan-binding loops.

**Figure 3.**
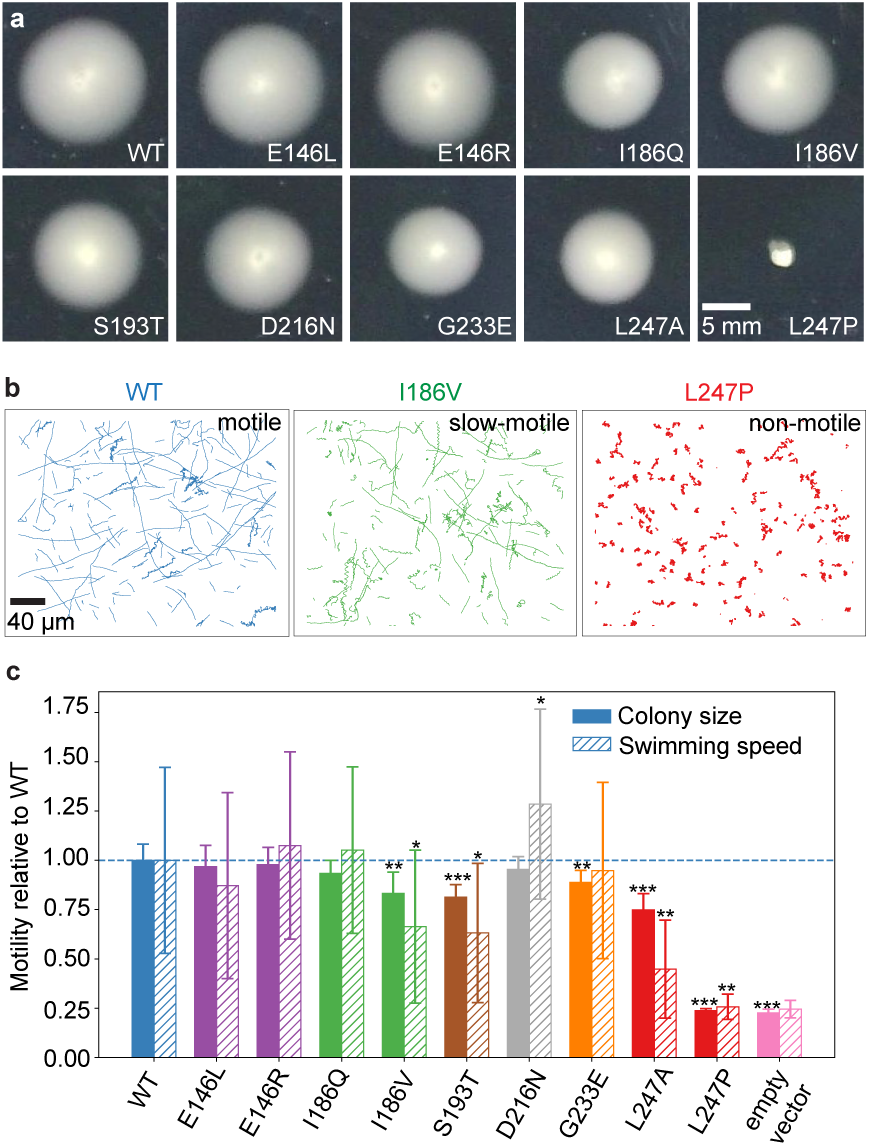
Cells expressing MotB stator mutants exhibit distinct motility phenotypes compared to WT. **a** Stator mutations alter swimming motility in soft agar. Representative colonies of motB stator mutants expressed from the pBAD33 plasmid in 0.25% soft agar with 0.02% arabinose show variable expansion sizes, reflecting differences in swimming motility. **b** Representative swimming trajectories of *E. coli* cells expressing WT, I186V, and L247P MotB stators. Faster swimmers exhibit straighter tracks, whereas slower swimmers show more curved trajectories. **c** Motility trends in soft agar and liquid are consistent across MotB mutants. Solid bars show normalized soft-agar colony expansion (square root of colony area), and hashed bars show normalized swimming speeds in liquid. Both soft agar and liquid swimming data were normalized to WT *motAB* expressed from pBAD33 at the same induction level. Error bars indicate standard deviation. Asterisks denote statistical significance relative to WT (*p* < 0.05, *p* < 0.01, *p* < 0.001; Student’s t-test for soft agar, Welch’s t-test for liquid). A t-test was not performed for the empty-vector control.

Highly motile cells (e.g., WT MotB) produced predominantly straight, fast trajectories, whereas slower variants (e.g., I186V) displayed curlier, slower tracks (Fig. 3b). As expected, the nonmotile L247P mutant exhibited only short, highly curved trajectories attributable to Brownian motion. Substitutions at the two rigid sites that showed the strongest co-evolution and dynamic coupling with the loops yielded contrasting phenotypes: cells expressing S193T MotB swam more slowly than WT and formed smaller soft agar colonies, whereas D216N MotB expressing cells swam faster than WT in liquid though they generated WT-sized colonies in soft agar. Mutations at the two flexible sites—E146 and I186, the most distal residues dynamically coupled to the loops—also yielded divergent results. While charge-altering mutations at E146 (E146R and E146L) had little effect on motility, a charge conserving substitution I186V reduced swimming speed. Surprisingly, the more drastic hydrophobic-to-polar substitution I186Q did not impair motility. Overall, almost all mutant stators remained functional but exhibited mutation-specific alterations in swimming speed, demonstrating that distal yet dynamically coupled sites within MotB’s periplasmic domain can modulate swimming motility *in vivo*.

### Stator expression titration reveals mutation-specific swimming-speed responses

Because swimming speed depends on both the number of stators assembled onto the motor and the torque produced by each stator, varying stator expression provides a way to test how each mutation affects stator function across different assembly conditions. We therefore measured swimming speeds across a five-order-of-magnitude range of stator expression (2 × 10^−5^%–2% Arabinose, w/v).

As stator expression increases, more stators assemble onto the motor (Fig. 4a), leading to higher swimming speeds until the motor reaches its maximum stator capacity (Reid et al., 2006). If a mutation weakens stator anchoring, the motor may fail to reach full stator occupancy even at high expression levels. Alternatively, if a mutation reduces the torque generated by each stator, the motor will exhibit a lower maximal swimming speed despite full occupancy. In both cases of reduced anchoring or reduced torque, mutants are expected to swim more slowly than WT across the entire range of stator expression (Fig. 4a).

**Figure 4.**
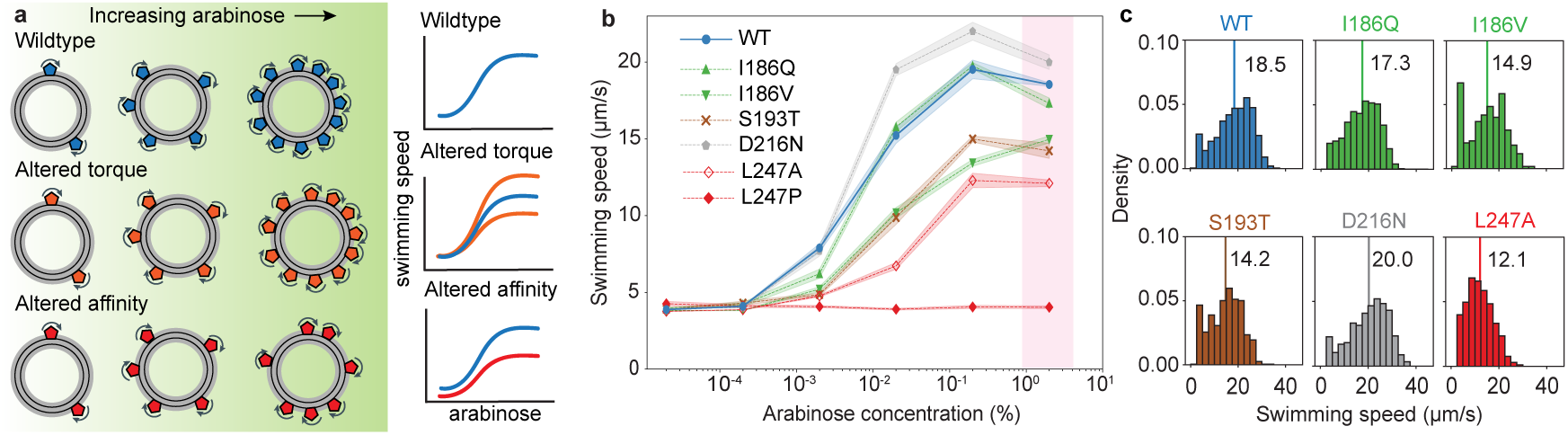
Expression-dependent swimming measurements reveal mutation-specific changes in stator function. **a** Top row: Increasing stator expression increases the number of motor-bound stators and swimming speed until saturation. Middle row: Altered torque production leads to different maximal swimming speeds at high stator expression. Bottom row: Mutations that reduce stator binding affinity limit stator occupancy and prevent full saturation even at high expression. **b** Most mutants exhibit swimming-speed trends distinct from WT across all arabinose concentrations, indicating altered stator function. Mean swimming speeds of *E. coli* expressing WT or mutant motB are shown; shaded regions indicate the standard error. The pink shaded area highlights swimming speeds at saturating arabinose concentrations. **c** Swimming-speed distributions at 2% arabinose reveal mutation-dependent population differences. Relative to WT, L247A and D216N show left- and right-shifted distributions, respectively, consistent with altered torque production, whereas I186V and S193T exhibit prominent low-speed peaks reflecting a large fraction of non-motile cells, consistent with altered stator anchoring. Vertical lines denote mean swimming speeds.

Swimming speeds increased with stator expression for all mutants except the nonmotile L247P variant, which remained immotile across all conditions (Fig. 4b). Several mutants exhibited consistently reduced swimming speeds relative to WT across the full expression range. In particular, I186V, S193T, and L247A swam more slowly than WT at all arabinose concentrations and saturated at lower maximal speeds. In contrast, I186Q closely tracked WT across expression levels, whereas D216N swam faster than WT over most of the range.

To further examine differences among mutants at high expression, we compared swimming-speed distributions across entire populations at 2% arabinose (Fig. 4c). Distributions for I186V and S193T contained a substantial fraction of nonmotile and slow-moving cells, broadening their distributions and explaining the lower mean swimming speeds compared to WT. For L247A and D216N, the fraction of non-motile cells was similar to WT, but their swimming-speed distributions were shifted leftward and rightward, respectively, consistent with altered torque production. These population-level differences complement the expression-dependent trends and reveal distinct mutation-specific signatures in swimming behavior.

### Single motor measurements distinguish altered torque from altered stator dynamics

To directly assess whether mutations affected torque production by individual stators, we performed tethered-cell assays under high stator expression conditions (2% arabinose), where motors are expected to operate under high load and be saturated with stators (Fig. 5a). Under these conditions, differences in motor rotation frequency primarily reflect differences in torque output rather than stator number or binding dynamics.

**Figure 5.**
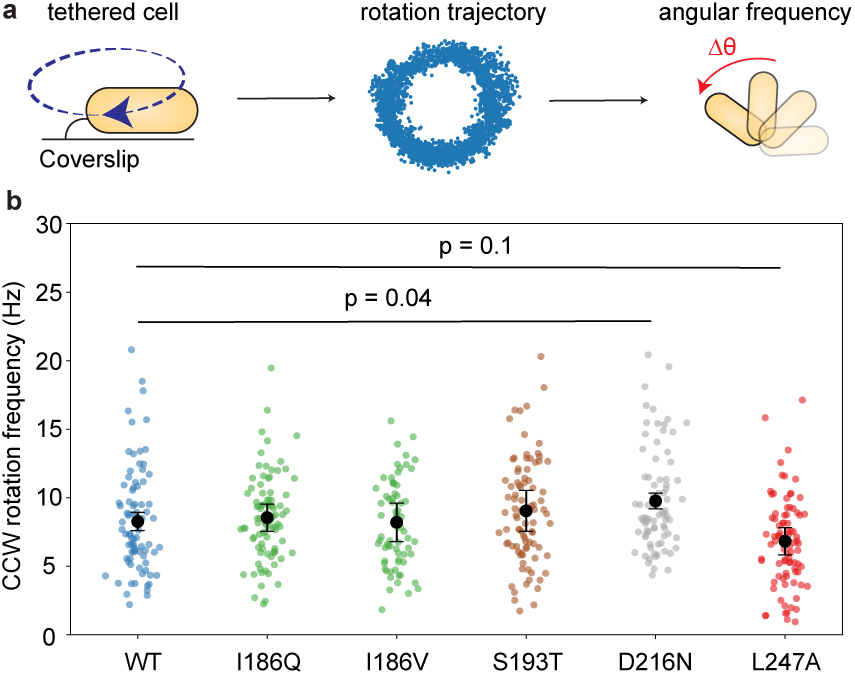
Single motor measurements distinguish altered torque production from altered stator anchoring. **a** Rotational frequency was determined by tracking the angular displacement, Δ*θ*, of tethered cells between consecutive frames. **b** Rotational frequencies of tethered cells under high load at saturating stator expression (2% arabinose) identify stator mutants with altered torque. Black dots and error bars denote mean frequency and standard deviation, respectively. D216N and L247A exhibit higher and lower rotation frequencies than WT, consistent with altered torque production, whereas I186Q, I186V, and S193T rotate at frequencies comparable to WT.

Motors driven by the L247A stators rotated more slowly than WT, whereas motors driven by the D216N stators rotated significantly faster, indicating reduced and enhanced torque production, respectively. In contrast, motors driven by I186Q, I186V, and S193T stators rotated at frequencies comparable to WT, suggesting that these mutations do not alter torque generation (Fig. 5b). Therefore, the observed differences in motility for these mutants (Fig. 3c) must be due to changes in stator dynamics under those conditions. Within the applied area cutoff (2-3 µm^2^), the distribution of cell sizes did not differ significantly across mutants (Fig. S5a), indicating that differences in rotation frequency were not attributable to cell size. Although the I186V mutant exhibited an increased clockwise (CW) bias relative to WT and other mutants (Fig. S5b), this did not affect our analysis because rotation frequencies were computed using only counterclockwise (CCW) intervals.

These single-motor measurements complement and explain the swimming trends observed in the arabinose titration experiments (Fig. 4b). The low- and high-torque mutants, L247A and D216N, were also the slowest and fastest swimmers, respectively, across expression levels. In contrast, mutants I186V and S193T saturated at lower swimming speeds despite exhibiting WT-like rotation frequencies in tethered-cell assays, indicating that their reduced motility arises from altered stator dynamics rather than impaired torque production. This distinction is further supported by swimming-speed distributions at high stator expression: I186V and S193T showed broadened distributions with an increased fraction of slow-moving cells, whereas L247A exhibited a uniformly left-shifted distribution consistent with reduced torque output (Fig. 4c). Together, these results demonstrate that mutations distal to the peptidoglycan-binding interface affect stator dynamics, whereas mutations at or near the binding interface affect torque generation, revealing two distinct mechanisms by which MotB mutations modulate motor function.

### Loop flexibility of mutant stators predicts their swimming performance

To assess how mutations at coupled sites influence conformational dynamics of the MotB periplasmic domain, we performed all-atom molecular dynamics simulations for WT MotB and selected mutant variants. From these simulations, we computed dynamic flexibility index (DFI) profiles for each MotB variant, quantifying the relative flexibility of each residue in the context of the global protein network dynamics (see Materials and Methods), thereby capturing how loop motions are affected by distal mutations.

Analysis of dynamic flexibility profiles revealed mutation-dependent changes in the mobility of the periplasmic loops implicated in stator anchoring. While loop 3 was largely flexible across all variants, the dynamics of the other two loops varied substantially between mutants (Fig. S6). Even the distal mutations altered the collective flexibility profile of the loop region as a whole (Fig. S6). In particular, substitutions at position 186 produced contrasting outcomes (Fig. 6a). In the I186Q mutant, loop 2 became largely rigid, while loops 1 and 3 exhibited reduced and spatially heterogeneous flexibility compared to WT, which maintains intermediate flexibility across all three loops. In contrast, the I186V substitution increased flexibility in portions of loops 1 and 2 relative to WT. Together, these results demonstrate that mutations at a distal site can differentially modulate the dynamic profiles of the peptidoglycan-interacting loops.

**Figure 6.**
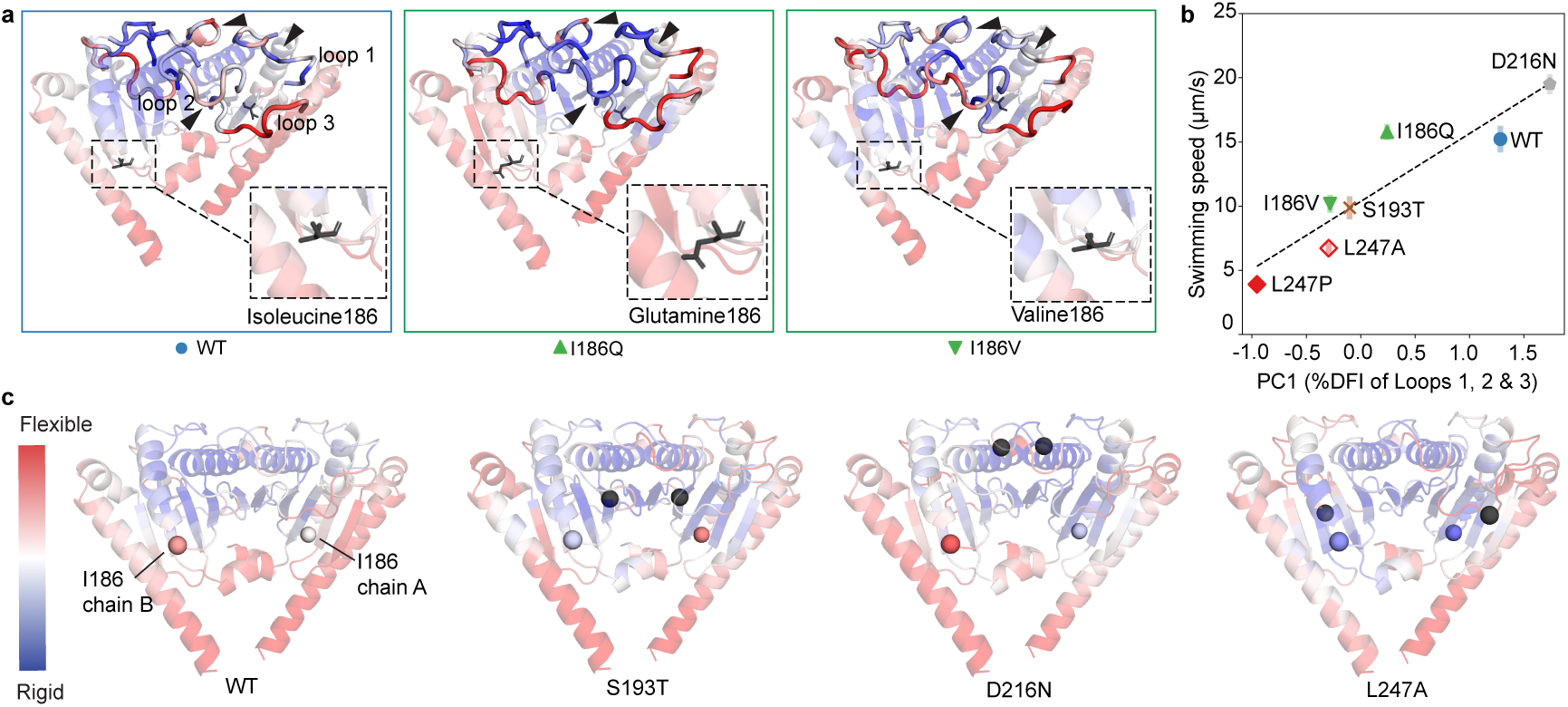
Loop flexibility links long-range coupling to stator performance. **a** Substitutions at position 186 have distinct effects on the flexibility profile of the peptidoglycan-interacting loops. Dynamic flexibility indices from MD simulations are shown, with flexible regions in red and rigid regions in blue. Insets highlight the residue substitutions at position 186, and black arrows mark regions with differential flexibility between variants. **b** Swimming performance of stator variants correlates strongly with the dominant mode of loop flexibility. Swimming speeds measured at 0.02% arabinose are plotted against the first principal component (PC1) of the loop DFI profiles (R = 0.917, p = 0.0036). Markers represent means and error bars indicate the standard error. **c** Mutations at other distally coupled sites also affect flexibility at position 186. In WT, residue 186 is predominantly flexible, whereas in the low-torque mutant L247A it becomes rigid in both chains. In contrast, in the other mutants, position 186 exhibits asymmetric flexibility across the dimer, remaining flexible in one chain and rigid in the other.

To quantify dominant patterns of loop dynamics, we performed principal component analysis, a dimensionality reduction method that extracts the dominant sources of variation across the DFI profiles of the three peptidoglycan interacting loops. Strikingly, the first principal component (PC1), which captured the largest variance in loop dynamics, showed a strong correlation with the experimentally measured swimming speeds (R = 0.917, p = 3.6 × 10^−3^) (Fig. 6b). Particularly, the non-swimming mutant L247P and the fastest swimming D216N mutant occupy the two extremes of the PCA axis. This relationship held across both distal mutations and mutations proximal to the anchoring interface, strongly suggesting that mutation-driven alterations in loop dynamics directly underlie changes in stator function and motor-level behavior.

Notably, mutations at other coupled sites also altered the flexibility profile at position 186 (Fig. 6c), indicating that this residue both controls the dynamics of distant regions, i.e., peptidoglycan-interacting loops, and is itself controlled by other distal residues. In the D216N and S193T mutants, residue I186 exhibited asymmetric changes across the MotB dimer, becoming more flexible in one subunit and more rigid in the other relative to WT. In contrast, the L247A mutation rigidified I186 in both subunits. Together, these observations indicate that position I186 both influences and responds to changes in loop dynamics, consistent with bidirectional dynamic coupling within the periplasmic domain.

Together, these results demonstrate that mutation-induced changes in MotB loop dynamics quantita-tively track swimming performance and identify dynamically sensitive residues that respond to both local and distal perturbations.

## DISCUSSION

In this study, we investigated how mechanical load is transmitted within the MotB periplasmic domain to regulate stator anchoring and motor performance. By combining coarse-grained elastic-network modeling, targeted mutagenesis, and quantitative motility measurements, we show that residues distant from the peptidoglycan-binding interface nonetheless exert strong control over stator behavior. Our results reveal that “long-range dynamic coupling” within MotB tunes stator function through two distinct mechanisms: modulation of stator anchoring dynamics and direct alteration of torque output. We demonstrate that variations in flexibility in peptidoglycan-binding loops quantitatively predict stator performance, providing a mechanistic link between structure, dynamics, and function. Together, these findings provide a mechanistic framework for understanding how mechanical information propagates through the stator to enable adaptive remodeling of the bacterial flagellar motor.

A key finding of this study is the presence of long-range coupling within the MotB periplasmic domain. Dynamic flexibility and coupling analyses identified the peptidoglycan- and P-ring-interacting loop 3 as a highly flexible element within the domain. These dynamic signatures align with previous observations of distinct open and closed conformations of these loops in the *H. pylori* MotB periplasmic domain (Reboul et al., 2011). Our results further indicate that loop motions are not locally determined but strongly linked to residues far from the anchoring interface, including sites across the dimer. Coevolutionary analysis independently supported these connections, revealing evolutionary coupling between dynamically linked residues despite separations that preclude direct contact. Together, these results indicate that MotB encodes intrinsic long-range communication pathways that transmit conformational and mechanical information across the periplasmic domain, suggesting that stator anchoring is regulated by the global dynamic architecture of MotB rather than by local interactions alone.

A striking outcome of perturbing these dynamically coupled sites through targeted mutagenesis is that distal mutations produce two qualitatively distinct effects on stator function. Some mutations primarily alter stator dynamics, leading to reduced or heterogeneous stator engagement without substantially affecting the torque generated by individual stators. In contrast, other mutations directly alter torque output. This distinction was revealed by combining expression-dependent swimming measurements with single-motor tethered-cell assays, which allowed anchoring-related effects to be separated from intrinsic torque generation. Notably, mutations at or near the anchoring interface (e.g., D216N and L247A) primarily altered torque output, consistent with local disruption of force transmission. In contrast, mutations at distal sites (e.g., I186V and S193T) affected stator dynamics without directly impairing torque generation, indicating that long-range coupling within MotB enables remote regulation of stator engagement.

Beyond these categorical distinctions, the correspondence between the dominant mode of variation in loop dynamics and swimming performance points to loop flexibility as a key control parameter for stator function. Both local and distal perturbations appear to influence motility by reshaping the conformational dynamics of the peptidoglycan-binding loops. In this context, residue I186 emerges as a central tuning site, capable of both responding to and propagating long-range perturbations within the periplasmic domain. Such tuning sites may allow stator behavior to be adjusted continuously without drastically affecting the core architecture of the motor.

These findings have direct implications for understanding load-dependent stator remodeling in the bacterial flagellar motor. Previous work has shown that stator units form catch bonds with the peptidoglycan, such that increased mechanical load stabilizes stator attachment by reducing unbinding rates Nord et al. (2017); Wadhwa et al. (2019, 2021). Our results suggest that this mechanosensitive behavior may be mediated, at least in part, by load-induced changes in the conformational ensemble of the MotB periplasmic domain (Haliloglu and Bahar, 2015; Nussinov, 2025). Mechanical load acting on an active stator must propagate through the stator from the MotA-MotB interface to the peptidoglycan-anchoring domain of MotB (Fig. 7a), providing a route by which load can tune the flexibility of the anchoring loops through long-range coupling (Fig. 7b). By tuning loop flexibility, MotB could regulate the lifetime of stator attachment at the peptidoglycan interface, providing a dynamic mechanism for stator recruitment and retention during motor remodeling (Fig. 7c).

**Figure 7.**
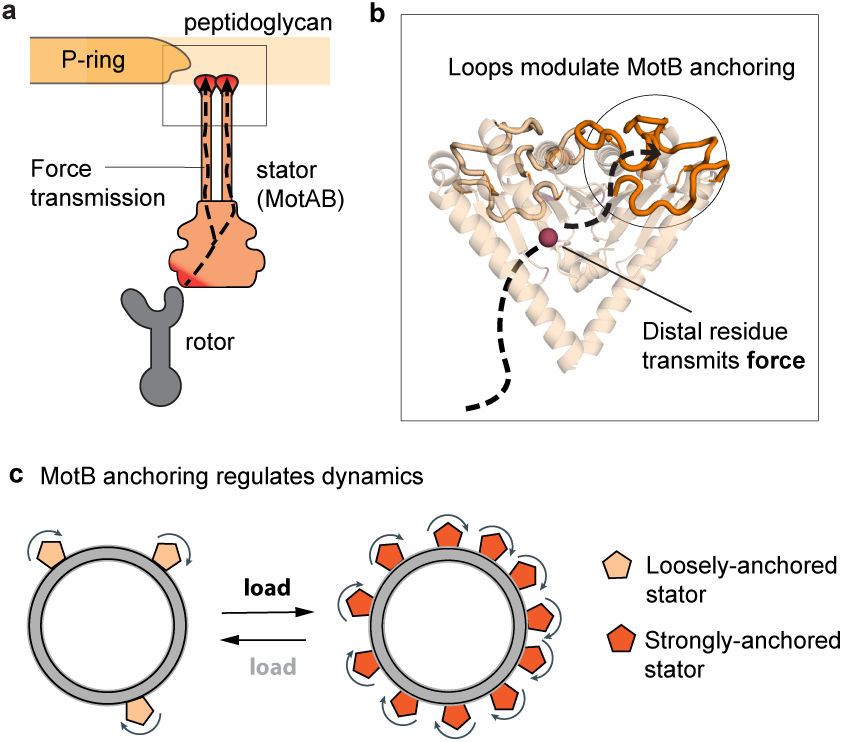
A conceptual model for dynamic allosteric regulation of stator anchoring by MotB. **a** Load applied at the MotA–rotor interface is transmitted through the stator to MotB’s periplasmic domain. **b** Distal residues within the MotB periplasmic domain transmit force to reshape the conformational ensemble of the anchoring loops via long-range coupling. **c** These dynamic changes regulate stator lifetime at the peptidoglycan interface and enable stator remodeling under varying mechanical load.

In summary, this work identifies MotB as a dynamically responsive module whose internal coupling enables mechanical information to be transmitted across length scales within the stator. Long-range coupling is a common feature of allosteric proteins (Monod et al., 1965; Süel et al., 2003; Reynolds et al., 2011), and our results suggest that bacterial stators exploit this principle to enable stable anchoring while simultaneously permitting dynamic regulation. More broadly, these findings illustrate how allosteric coupling can endow molecular complexes with the ability to sense and respond to mechanical load, a strategy that may be widely employed in other force-bearing protein assemblies.

## MATERIALS AND METHODS

### Modeling MotB periplasmic domain

We modeled the *E. coli* MotB periplasmic domain using SWISS-MODEL and selected three of the top ten predictions (models 01, 05 and 07), each corresponding to a disctinct crystal form. We compared these models with structural prediction of the MotB periplasmic domain from AlphaFold by superimposing them in PyMOL. The overall root mean squared deviation (RMSD) values between the AlphaFold prediction and model 01, model 05 and model 07 were 0.702, 0.749 and 0.732 respectively.

### Dynamic flexibility index (DFI)

The dynamic flexibility index (DFI) is a residue-specific metric that quantifies how resistant a protein position is to force perturbations. It measures the relative fluctuation response of a residue compared to the overall fluctuation response of the protein (Kumar et al., 2015; Larrimore et al., 2017). The method is based on linear response theory applied within either molecular dynamics (MD)–derived covariance matrices or coarse-grained elastic network models (ENMs).

To compute DFI, we perturb each residue sequentially with a small, random Brownian force and calculate the fluctuation response of all residues. The displacement response vector Δ*R* is obtained from the linear response relationship:

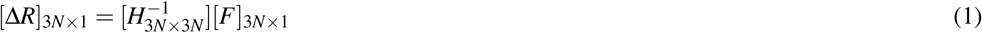

where *F* is the perturbing force vector, and *H* is the Hessian matrix of second derivatives of the potential energy. The inverse Hessian, *H*^−1^, corresponds to the covariance matrix that captures residue–residue couplings. Forces are applied in multiple directions to approximate isotropic fluctuations, and the averaged responses are used to obtain residue-level profiles.

The DFI of residue *i* is then defined as:

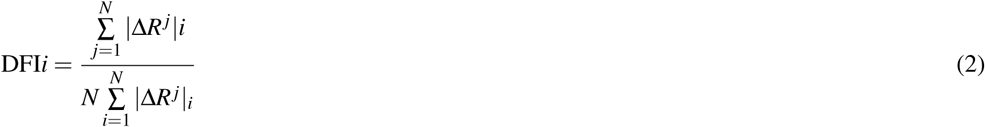

Here, |Δ*R^j^*|*_i_* is the magnitude of the displacement of residue *i* in response to perturbation at residue *j*, and the denominator normalizes by the total response across the protein. Thus, the DFI represents the fractional contribution of residue *i* to the global fluctuation response.

Residues with low DFI values (<0.2) act as hinge points within the interaction network. These sites typically exhibit limited local fluctuations yet serve as effective conduits for transmitting perturbations, thereby exerting control over collective motions. In contrast, high-DFI residues are dynamically more flexible and less constrained by network interactions. Prior studies have shown that mutations at low-DFI positions often disrupt communication pathways and lead to functional changes or disease associations (Campitelli et al., 2020, 2025). Substitutions at such sites frequently modify catalytic activity, ligand binding, or allosteric communication by shifting equilibrium dynamics.

In this study, covariance matrices used for DFI calculations were obtained from both MD simulations (for all variant and WT) and ENM (Fig. 2). The resulting DFI profiles were then analyzed using principal component analysis (PCA) to identify dominant dynamical trends associated with MotB function.

### Dynamic Coupling Index (DCI)

The dynamic coupling index (DCI) quantifies how strongly the fluctuation response of residue *i* is coupled to perturbations at a specific residue *j*, relative to its average response to perturbations at all residues in the protein. Thus, DCI compares a specific response to a global baseline response, highlighting preferential dynamic couplings between residues.

Formally, the DCI of residue *i* with respect to residue *j* is defined as:

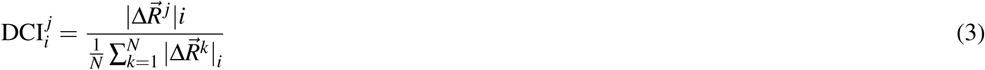

Here, 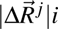 is the fluctuation response of residue *i* to perturbation at residue *j*, and the denominator represents the average fluctuation response of residue *i* to perturbations across all *N* residues. A value of 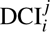 > 1 indicates that residue *i* is more strongly coupled to *j* than expected on average, whereas 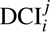 < 1 indicates weaker-than-average coupling.

DCI has been widely used to identify residues that mediate long-range communication, even when separated by large structural distances. For example, residues far from the active site can exhibit high DCI with catalytic residues, implicating their role in allosteric regulation (Campitelli and Ozkan, 2019; Campitelli et al., 2020). Beyond enzymes, DCI has been applied to uncover the role of distal sites in viral protein allostery, protein–RNA interactions, and viral capsid assembly (Hamilton et al., 2024; Ose et al., 2024).

Together with DFI, which identifies flexible and rigid communication hubs, DCI provides a complementary measure of residue–residue coupling that enables the detection of allosteric pathways and mechanistic insights into how mutations reshape protein dynamics.

### Co-evolution

The method to compute co-evolution relies on sequence information to build a matrix of structural contacts. We used the EVcouplings (Hopf et al., 2019) server to gather information on evolutionary couplings (ECs) within the MotB periplasmic domain. While these analyses are generally used to predict protein 3D structures as residue pairs with high ECs are expected to be in close proximity (Hopf et al., 2014), using our model structures we identified structurally distant residue pairs showing high evolutionary coupling. From the 12967 co-evolving pairs in the entire MotB sequence, we selected 328 pairs having a co-evolution probability equal to and above 80%. Because residue co-evolution is also an indicator of direct interactions in the folded protein (Hopf et al., 2012), we calculated distances between the C-*α* atoms of the co-evolving residue pairs in the folded structure of the MotB periplasmic domain to identify residue pairs that were spatially separated across different regions of the protein. Distant co-evolving residue pairs which also showed a high DCI value were found to be divided into three categories by further integrating information from the DFI analysis —1) flexible and co-evolutionarily coupled (E146 and I186), rigid and 2) co-evolutionarily coupled (S193 and D216) and 3) intermediate flexibility and co-evolutionarily coupled (G233).

### Bacterial Strains, Plasmids, and Cultures

All strains used here were derivatives of the *E. coli* strain MG1655. Strain NW165 was deleted for *motAB* using the lambda red scar-less recombination method (Murphy and Campellone, 2003) with a modified counter-selection system (Kim et al., 2020). To investigate the motility phenotype of mutant stators, plasmids expressing these mutants were constructed by first cloning *motAB* in pBAD33 using the NEBuilder HiFi DNA Assembly cloning kit. The pBAD33-*motAB* plasmid was then used as a template for the site-directed mutagenesis of MotB using Agilent’s Quick Change Kit. The resulting plasmids expressing *motA* along with one of the *motB* variants were then transformed into NW165. Strains for single motor measurements were transformed with the plasmid pFD313 which expresses a sticky filament.

Cells were grown by diluting an overnight culture 1:100 in TB (10 g/L tryptone, 5 g/L NaCl) supplemented with 25 µg/mL chloramphenicol, followed by incubation at 37^◦^C with shaking at 200 rpm to OD_600_ = 0.3. Cells were then induced with 2 × 10^−5^ − 2% arabinose until OD_600_ reached a range of 0.4–0.6.

### Soft agar assay

TB soft agar was prepared by dissolving 10 g/L tryptone, 5 g/L NaCl and 2.5 g/L of Difco agar in Milli-Q water and autoclaved. 25 mL of soft agar was poured into standard petri plates and supplemented with filter-sterilized arabinose at a final concentration of 0.02%. A well isolated bacterial colony was picked with a 20 *µ*L pipette tip and stabbed till halfway in the solidified soft agar. Plates were incubated at 30°C for 14-15 h. Plates were imaged using custom-built imaging setup, and motility halo sizes were measured using an in-house written script in Python. A total of 8 replicates were performed for each strain.

### Swimming speed measurements

Cells were harvested from cultures grown to OD_600_ = 0.4-0.6, by centrifugation at 1500 g for 7 min. The pellet was re-suspended in a motility buffer (10 mM potassium phosphate, 0.1 mM EDTA, 10 mM sodium lactate, 1mM methionine and 66.7 mM NaCl, pH 7.4). The cell suspension was diluted 50X (OD_600_∼0.01), introduced into a 65 µL geneframe (Thermo Fisher Scientific), and sealed with a glass cover slip. 1 minute long recordings of freely swimming cells were made using a 20X objective at 20 frames per second using ThorLabs (model DCC1545M) camera. Cell tracking and speed measurements of individual cells were performed using Trackmate in ImageJ (Tinevez et al., 2017; Schindelin et al., 2012). Average swimming speeds were calculated using TrackMate’s SimpleSparseLAPTracker algorithm (Jaqaman et al., 2008) which is suitable for particles undergoing Brownian motion. We only used tracks longer than 15 frames to avoid noise from spurious tracks. Finally, we converted the pixel/frame speed values from TrackMate to *µ*m/s by multiplying with a conversion factor obtained with a stage micrometer.

### Single Motor Measurements

3 mL of cells were harvested from cultures grown to OD_600_ = 0.4–0.6 by centrifugation at 1,500 × g for 7 min. The pellet was washed once with motility buffer (10 mM potassium phosphate, 0.1 mM EDTA, 10 mM sodium lactate, 1 mM methionine, and 66.7 mM NaCl, pH 7.4), and the final pellet was up-concentrated by re-suspending in 1 mL of the same buffer. To shear the flagellar filaments, the suspension was passed between two syringes through 10 cm of 23-gauge tubing 90 times. Cells were pelleted and resuspended a second time, then introduced into a flow chamber formed by a coverslip and glass slide separated by double-sided tape, and incubated at room temperature for 10 min. Excess, untethered cells were removed by washing with motility buffer prior to imaging. Tethered cells were recorded for 1 min using a 40× objective at 100 frames per second with a FLIR Blackfly S camera.

Image analysis was performed using custom Python scripts. To minimize selection bias, tethered cells were identified using an automated cropping procedure. Cells were selected and cropped if they exhibited sustained movement, permitting pauses of up to 10 s. A subsequent manual screening step retained only those cells tethered by a single pole, ensuring that the mechanical load experienced across the dataset remained comparable. Images were binarized using Otsu’s method, and the resulting segmented objects were used to determine the center of rotation and to compute rotational frequency. Cells with a detected area outside of 2-3 µm^2^ were removed from the analysis so that the load of the cell body would be comparable.

### Molecular Dynamics Simulations

Molecular dynamics (MD) simulations were performed using the AMBER 20 software package to investigate the dynamic properties of WT MotB and selected mutants. The starting structural model was obtained from a previously described SWISS-MODEL homology model. Point mutations were introduced in PyMOL, and the ff14SB force field was used for protein parameterization. Each system was solvated in a rectangular box of TIP3P water molecules, ensuring a minimum distance of 16 Å between any protein atom and the box boundaries. Charge neutrality was achieved by adding Na^+^ and Cl^−^ counterions.

The simulation protocol consisted of three main stages: energy minimization, equilibration, and production. Energy minimization was carried out in two steps. First, 100,000 cycles of steepest descent minimization were applied with positional restraints on protein and solvent heavy atoms, allowing relaxation of the water molecules. This was followed by unrestrained minimization, during which the SHAKE algorithm was applied to constrain covalent bonds involving hydrogens. After minimization, the system was gradually heated from 0 K to 300 K over 50,000 steps with a 2 fs timestep, using harmonic restraints on protein heavy atoms. Hydrogen atom constraints were maintained throughout the heating stage. Equilibration was then performed under constant pres-sure and temperature (NPT) conditions to stabilize both system density and pressure.

Production simulations were conducted under NPT ensemble conditions at 300 K and 1 bar. Temperature was regulated using a Langevin thermostat with a collision frequency of 1.0 ps^−1^, and pressure was controlled using a Berendsen barostat. A timestep of 2 fs was used, with SHAKE applied to constrain bonds involving hydrogens. Each system, including the WT and all selected mutants, was simulated for 2 *µ*s of production dynamics, providing sufficiently long trajectories for comparative analysis of slow conformational changes. To ensure adequate sampling, trajectory convergence was carefully evaluated following established methods (Sawle and Ghosh, 2016; Kazan et al., 2023).

Finally, we applied principal component analysis (PCA) to DFI from MD simulations to segregate the MotB mutants by dynamics. This approach enables dimensionality-reduction by a linear combination of original variables into principal components (PCs) which capture the largest variability from the data (Jolliffe, 2005).

## AUTHOR CONTRIBUTIONS

S.A.S., S.B.O., and N.W. designed the research; N.W. and S.B.O. oversaw the research; S.A.S. performed experiments; S.A.S., B.M.W. and I.C.K. analyzed data; I.C.K. performed MD simulations; S.A.S., B.M.W., S.B.O., and N.W. wrote the manuscript.

## FUNDING

This work is supported by the National Institute of General Medical Sciences of the National Institutes of Health (Grant R01GM147635 to S.B.O and Grant R00GM134124 to N.W), Gordon and Betty Moore Foundation (AWD0003443 to S.B.O), and the Arizona Biomedical Research Centre (Award RFGA2023-008-14 to N.W).

## ACKNOWLEDGMENTS

We thank Karen Fahrner and Carolina Gogerty for helpful suggestions on strain construction, Keichi Namba and Mayra Garcia-Alcala for sharing plasmids, and Ngoc Huynh and Nikhil Ramesh for sharing data visualization scripts. We are also grateful to all members of the Wadhwa lab for their insightful discussions and feedback throughout this study.

## CONFLICTS OF INTEREST

The authors declare no conflict of interest.

## Notes

### Competing Interest Statement

The authors have declared no competing interest.

### Summary of Updates

Added new data, updated text and figures.

