## Supplementary Information for "Long-range coupling regulates stator dynamics in the bacterial flagellar motor"

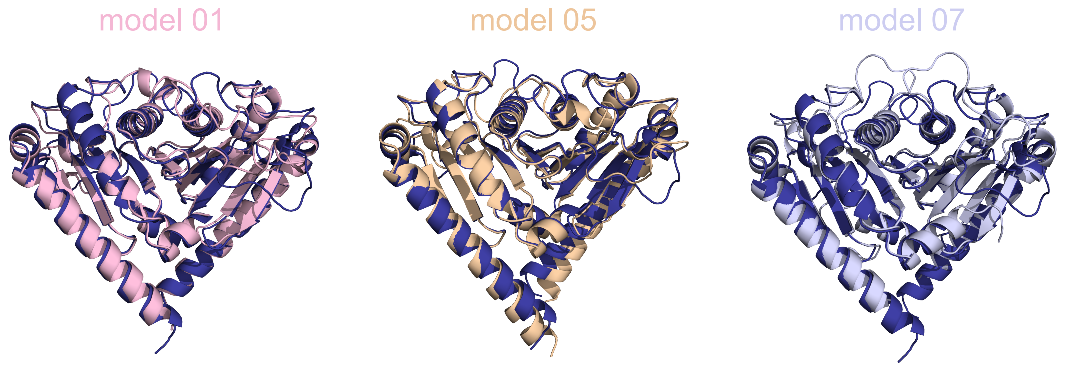


**Supplementary Figure S1. Superimposition of *E. coli* MotB models:** Structural predictions from SWISS-MODEL compare well with the prediction from AlphaFold3 (dark blue). Root Mean Squared Deviation (RMSD) values were computed by aligning structures in PyMOL. RMSD of AlphaFold structure with model 01 = 0.702, with model 05 = 0.749 and with model 07 = 0.732.

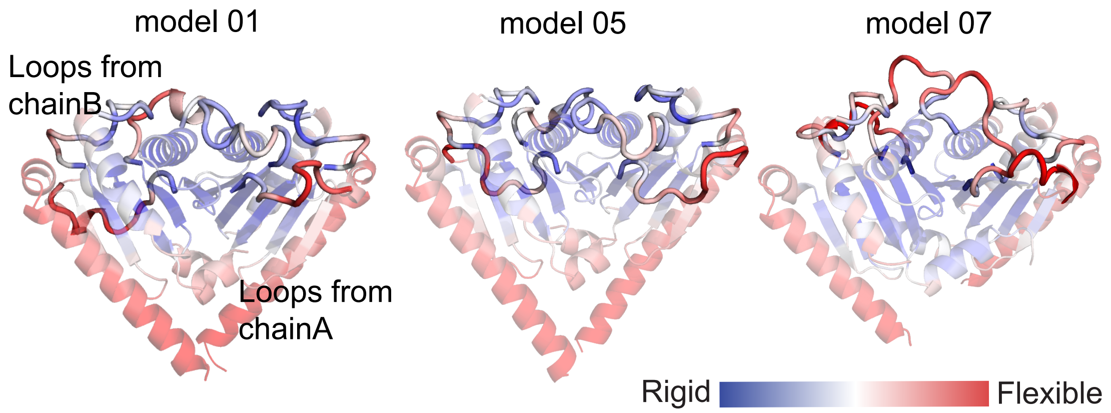


**Supplementary Figure S2. Dynamic profiles of all MotB periplasmic domain models.** Flexible (red) and rigid (blue) regions are represented. Peptidoglycan binding loops 1, 2 and 3 from both the chains A and B are shown as thicker cartoon representation.

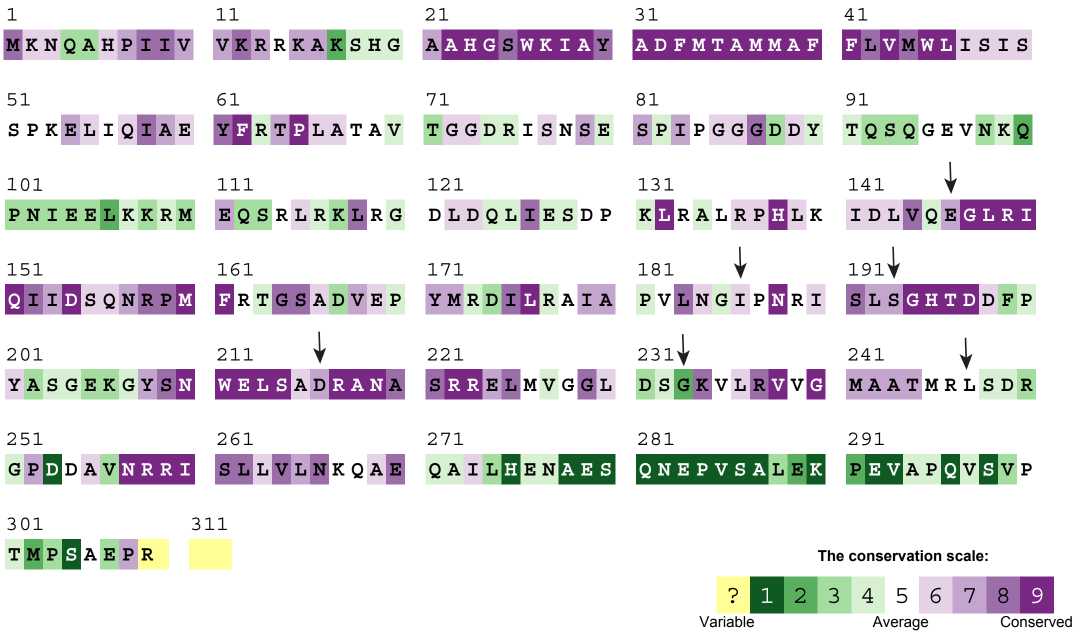


**Supplementary Figure S3. Sequence conservation of MotB gene from Consurf.** Arrows indicate residue positions that were chosen for making mutations.

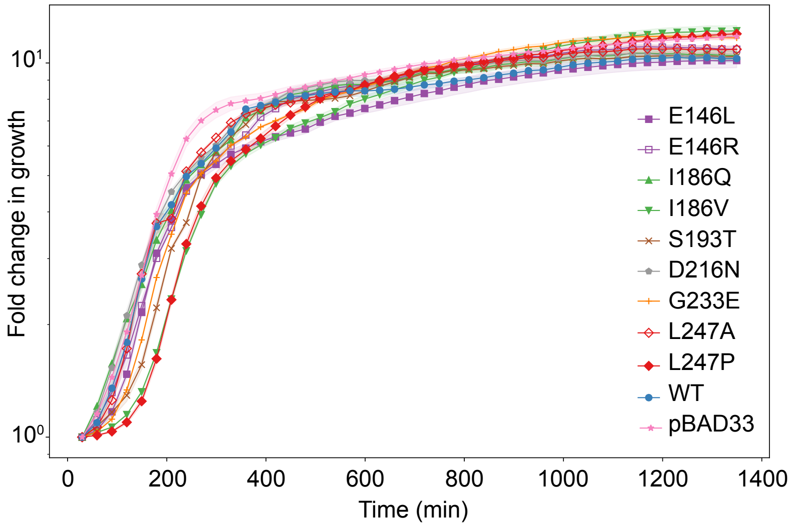


**Supplementary Figure S4. Growth curve of MotB mutants in LB supplemented with 0.02% arabinose.** The y-axis indicates fold change in growth for all strains monitored over 24 hours at 37°C. Optical density values were measured at a wavelength of 600 nm and normalized by the first reading. Markers indicate average from four replicate wells of a 96-well plate and the shaded region indicates the standard deviation.

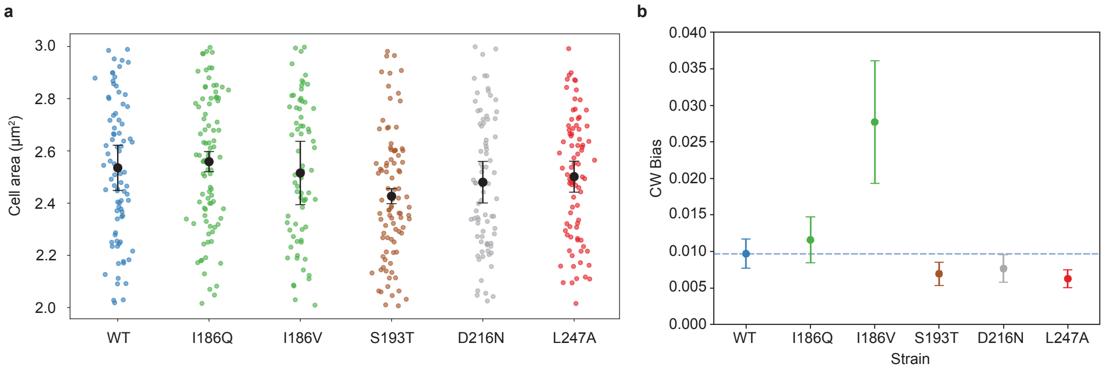


**Supplementary Figure S5. Additional statistics for tethered cell assays. (a)** Detected area of cell body after thresholding. For consistency, only cells with an area between 2 and 3 $\mu m^2$ were included in analysis. Black dots and error bars represent the mean and standard deviation. **(b)** Average clockwise bias of tethered cells, with error bars representing standard error of the mean. Dashed line shows WT CW bias for ease of comparison. CW bias is defined as the number of frames the cell rotates CW (negative frequency) divided by the total number of frames in a recording.

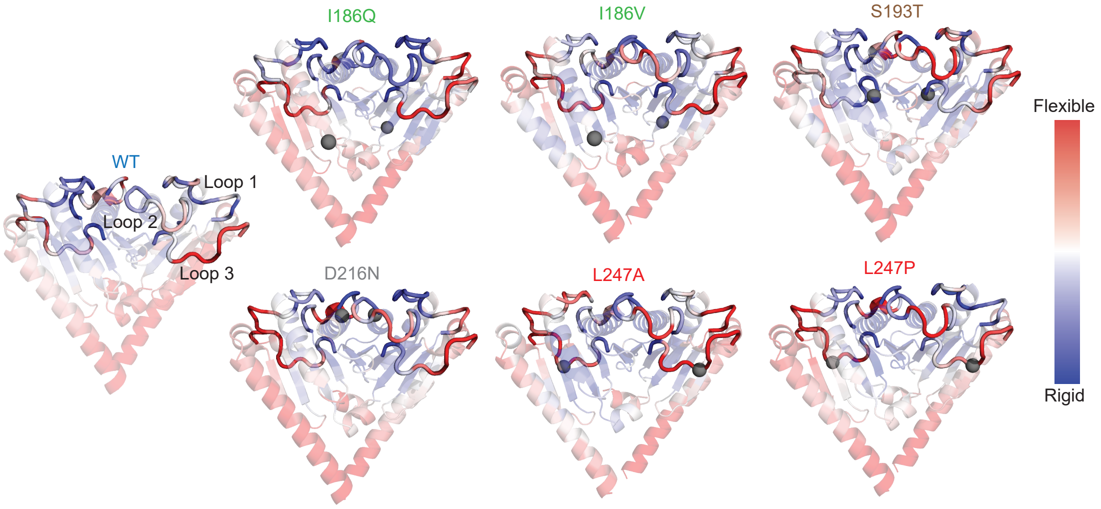


**Supplementary Figure S6. DFI values extracted from MD trajectories** Flexible (red) and rigid (blue) regions are shown. Mutation sites are represented as gray spheres. Mutations at distal sites affect flexibility of peptidoglycan interacting loops.

**Supplementary Table S1.** Dynamic flexibility (DFI), inter-chain dynamic coupling (DCI), and evolutionary coupling (EC) values for residues coupled with peptidoglycan interacting loops of MotB. Distances are given between C-$\alpha$ atoms.

| distal residue $(i)$ | pctDFI$_i$ | pctDCI$_{i-loop3}$ | loop residue $(j)$ | EC | Intra-chain (Å) | Inter-chain (Å) |
|---|---|---|---|---|---|---|
| 146 | 0.72 | 0.88 | 158 | 0.54 | 29.1 | 38.3 |
| 186 | 0.62 | 0.79 | 159 | 0.57 | 29.3 | 37.9 |
| 193 | 0.08 | 0.86 | 245 | 0.99 | 10.8 | 18.2 |
| 216 | 0.13 | 0.80 | 209 | 0.96 | 10.6 | 11.9 |
| 233 | 0.42 | 0.99 | 241 | 0.67 | 28.0 | 8.5 |
